# Environmental thresholds in the functional mycobiome of global drylands

**DOI:** 10.1101/2022.03.01.482255

**Authors:** Eleonora Egidi, Manuel Delgado-Baquerizo, Miguel Berdugo, Emilio Guirado, Davide Albanese, Brajesh K. Singh, Claudia Coleine

## Abstract

Fungi are major drivers of ecosystem functions. Increases in aridity are known to negatively impact fungal communities in dryland ecosystems globally, however, much less is known on the potential influence of other environmental drivers. To fill this knowledge gap, we reanalyzed fungal data from 912 soil samples, providing the largest and most complete fungal community dataset from global drylands. We used machine learning tools to examine geographical patterns in community composition and spatial, edaphic, and climatic factors driving them. Further, we determined critical thresholds of community turnover along those gradients. Our analysis identifies UV index, climate seasonality, and sand content as the most important environmental predictors of community shifts, harbouring greatest association with the richness of putative plant pathogens and saprobes. Important nonlinear relationships existed with each of these fungal guilds, with increases in UV and temperature seasonality above 7.5 and 900 SD, respectively, being associated with an increased probability of plant pathogens and unspecified saprotrophs occurrence. Conversely, these environmental parameters had a negative relationship with litter and soil saprotrophs richness. Consequently, these functional groups might be differentially sensitive to environmental changes, which might result in an inevitable disturbance of current plant-soil dynamics in drylands.

## Introduction

Drylands are the largest terrestrial biome (covering about 41% of the land surface and supporting 40% and 35% of the global population and global diversity, respectively) and are expected to expand further up to 56% by the end of the century ^1^. Drylands play key roles in regulating the global carbon, nitrogen and water cycles, and are thus fundamental for sustaining life on Earth ^2^. Due to their extreme temperatures, low and variable rainfall, and low soil fertility, drylands are particularly sensitive to changes in climate that lead to increased aridity (i.e, precipitation/potential evapotranspiration) ^3^.

Fungi are paramount components and drive critical ecosystem services in drylands, contributing to the formation of fertile islands ^4^, nutrient cycling and climate regulation ^5^, with a major role in dryland primary production ^6^ and pedogenesis ^7^. Key fungal groups include pathogens, mutualistic symbionts of both plants and animals, lichenized fungi, as well as soil and litter saprobes. Most previous studies on the biogeography and ecological attributes of fungal communities in dry systems have focussed on the role of aridity, given its role as a key driver of dryland ecology ^3,8–9^. However, other environmental factors could be potentially important in predicting fungal diversity and distributions in global drylands. For example, solar UV radiation is a primary driver of litter and soil organic carbon decomposition and plant growth in many arid and semi-arid ecosystems ^10,11^, suggesting a potential major contribution to the occurrence of decomposers and plant-associated fungi ^12^. Similarly, temperature and precipitation seasonality regulate plant cover dynamics and productivity in arid systems ^13^, which in turn can influence soil physical attributes important for saprotrophs, pathogen and mutualists distribution, such as soil moisture, pH, structure or carbon content ^14,15^.

Despite the possible centrality of multiple environmental variables in determining the spatial distributions of important fungal groups, their relative contribution to fungal biogeographical patterns remains largely unexplored at larger scales ^16^. Given the ecological and economic significance of drylands, and the global role of fungi in regulating their functions, it is critical to identify the environmental factors associated with distributions of fungal communities, and most importantly, to test whether the dependence of fungi on those drivers are linear or non-linear. The latter is important because non-linear associations between fungal distributions and environmental predictors, may signal particular environmental scenarios of exacerbated sensitivity (i.e., thresholds). Such abrupt shifts can mark regime shifts with potential implications for ecosystem functioning, which should be monitored and managed closely if we want to prevent changes of high magnitudes in the functional roles of fungi in a context of climate change ^17^. A better understanding of the forces shaping the global biogeography of dryland soil fungi can improve our ability to predict their fate under global change, and therefore inform future conservation and management policies.

Towards this aim, we conducted a meta-analysis of multiple datasets from different dryland biogeographical regions, merging sequencing data from a wide range of ecosystem and climates (i.e. hot, temperate, and cold drylands) and encompassing a representative plethora of all dryland sub-types (i.e. from hyper arid to dry-sub humid). We generated an unprecedented database of 1,473 fungal genera from a total of 912 individual topsoil samples (top 5-10cm) from all continents, including Antarctica. We examined geographical patterns in fungal assemblages and the main environmental (spatial, edaphic, and climatic) factors driving them as well as to establish where, along a range of environmental pressure gradients, important changes in community composition occur to identify critical thresholds along those gradients.

## Results

### General description of the dataset

Our dataset represents the largest extant fungal community dataset from drylands. Compared to previous large-scale studies focused on fungal diversity in drylands, our survey encompasses all continents, including Antarctica, and spans all dryland subtypes (defined by their aridity ranges), from hyperarid (AI ≤ 0.05, n = 38), to arid (0.05 < AI ≤ 0.2, n = 274), semiarid (0.2 < AI ≤ 0.5, n = 355), and dry sub-humid (0.5 < AI < 0.65, *n* = 265) regions of the world. Samples were distributed across cold (*n* = 378), temperate (*n* = 458) and hot drylands (*n* = 71) (Supplementary Information, Figure S1).

Of the 1,473 genera of fungi retrieved after bioinformatics analysis, 60% belonged to Ascomycota, 33% to Basidiomycota and 2.6% to Glomeromycota and 2% to Zygomycota (Supplementary Information, Figure S2). Out of the 66% (986) of taxa, 34.9% were saprotrophs (including 11.5% wood saprotroph, 9% litter saprotrophs, and 8% soil saprotrophs), 13% plant pathogens, 8% endophytic-mycorrhizal (5% ectomycorrhizal, 1% arbuscular-mycorrhizal, and 2% root-foliar endophytes/epiphytes), and 5% were lichenized (Supplementary Information, Figure S3). Plant pathogens were mostly dominated by Dothideomycetes (37.6%) and Leotiomycetes (10%), while ectomycorrhizal fungi, wood, litter and soil saprotrophs were dominated by Agaricomycetes (71, 60.5, 32 and 30%, respectively) (Supplementary Information, Figure S4).

### Environmental drivers of functional composition

The relative importance of spatial, edaphic, and climatic variables in predicting the composition of the main fungal ecological groups was determined using Gradient Forest (GF) methods, which identified the major determinants of community composition of fungi. The total model prediction performance from the GF analysis (i.e., the proportion of variance explained in a random forest) was averaged across the suite of environmental variables from the most common guilds (i.e., among those occurring in at least 10% of the samples), and ranged from 0.01 to 0.12 (R-squared; Figure 1A). Greatest global community turnover was associated with the spatial variables (PCNM1 and PCNM1 eigenvector-based vectors, maximum cumulative importance: 0.12 and 0.08, respectively), followed closely by UV index (UV), with a maximum value above 0.07. Importance in relation to other environmental predictors was highest (> 0.04) for diurnal temperature range (DTR), sand and temperature seasonality (TSEA), while mean annual temperature (MAT), precipitation seasonality (PSEA), pH, aridity index (AI) and soil organic content (SOC) had the lowest importance values (0.02-0.04) (Figure 1A).

**Figure 1.**
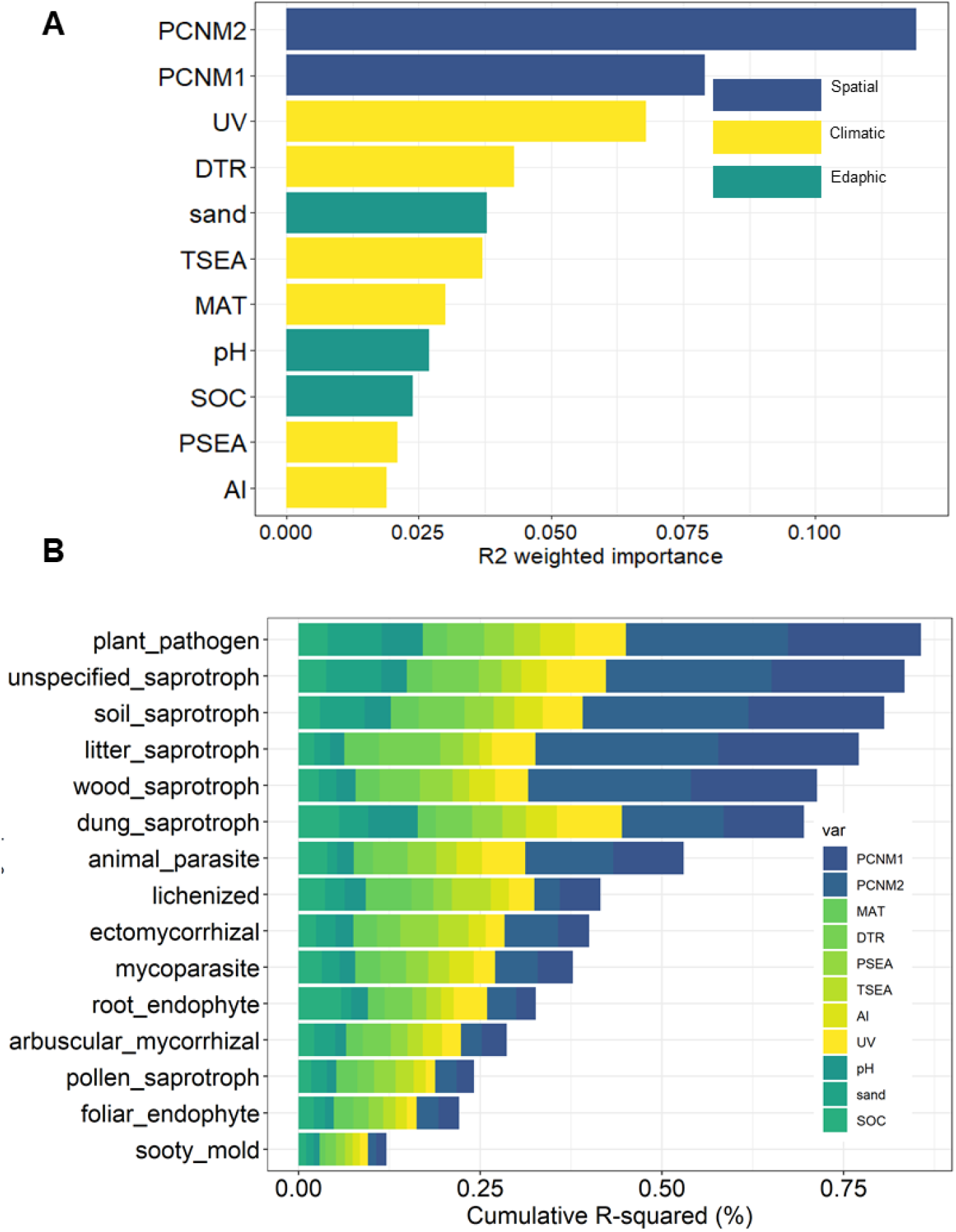
Environmental predictors of dryland fungal community composition. **(A)** Relative importance, R-squared (R^2^), of each environmental predictor included in the gradient forest analysis. **(B)** Contribution (0 to 1) of climatic, soil and spatial categories to the variation explained by the complete gradient forest model for the 15 ecological groups for which significant predictive power was established (R-squared >0).

Then, for each ecological group, we identified the most important predictors of changes in their abundance along spatial, climatic, and edaphic gradients. The cumulative model prediction performance of the guilds for which significant predictive power was established (R-squared >0) had a range of 0.11–0.86 (R-squared), with the highest model performance (>0.70) recorded for plant pathogens and soil, litter, wood and unspecified saprotrophs (Figure 1B). The predictive power of PCNM1 and PCNM2 was strongest for these fungi relative to other ecological groups (R-Squared values > 0.10; Supplementary Information, Figure S5). However, plant pathogen and unspecified saprotroph richness were also strongly predicted by UV radiation and sand content (R-Squared values 0.07-0.08, respectively), while DTR was the single most important climate predictor of litter saprotroph richness, followed by UV (R-Squared values = 0.08 and 0.06, respectively). DTR was also important in predicting soil saprotrophs distributions, together with sand content (R-Squared values = 0.06 for both). Conversely, PSEA and TSEA were the strongest predictors of ectomycorrhizal fungi (R-Squared values=0.04 and 0.05, respectively), with TSEA also strongly associating with lichenised fungi (R-Squared values=0.05; Supplementary Information, Figure S5).

### Detection of thresholds

Frequency histograms and density plots of the values used by the classification trees for splits (i.e. the split density plots in Figure 2A) were utilised to quantify thresholds at whole community scales (i.e., where important community changes occur) along the most predictive environmental gradients. UV index harboured a major threshold at values > 7, where most of the data occurred (Figure 2A). This threshold corresponded to a shift in the proportion of most ecological groups, including plant pathogens, litter and dung saprotrophs, as indicated by the steep slope in the cumulative plots (Figure 2B). Along the two secondly most important climatic variables (DTR and TSEA), we observed multiple subsequent strong splits. The fungal community showed a first response with mean diurnal temperature range > 8 °C, and then a second threshold with mean diurnal range > 14 °C, the latter mainly corresponding to shift in proportion of a range of saprotrophic fungi (i.e, litter, soil and unspecified saprotrophs); shifts in lichenized and ectomycorrhizal fungi, as well as animal and plant pathogens and dung saprotrophs, was recorded with a variation of monthly temperature averages > 1,000 SD and > 500 SD (Figure 2A and B). Finally, important community changes related to soil edaphic features were brought about by sand content between 45-60%, with shifts in plant pathogens and a range of saprotrophs (unspecified, dung and soil; Figure 2A and B).

**Figure 2:**
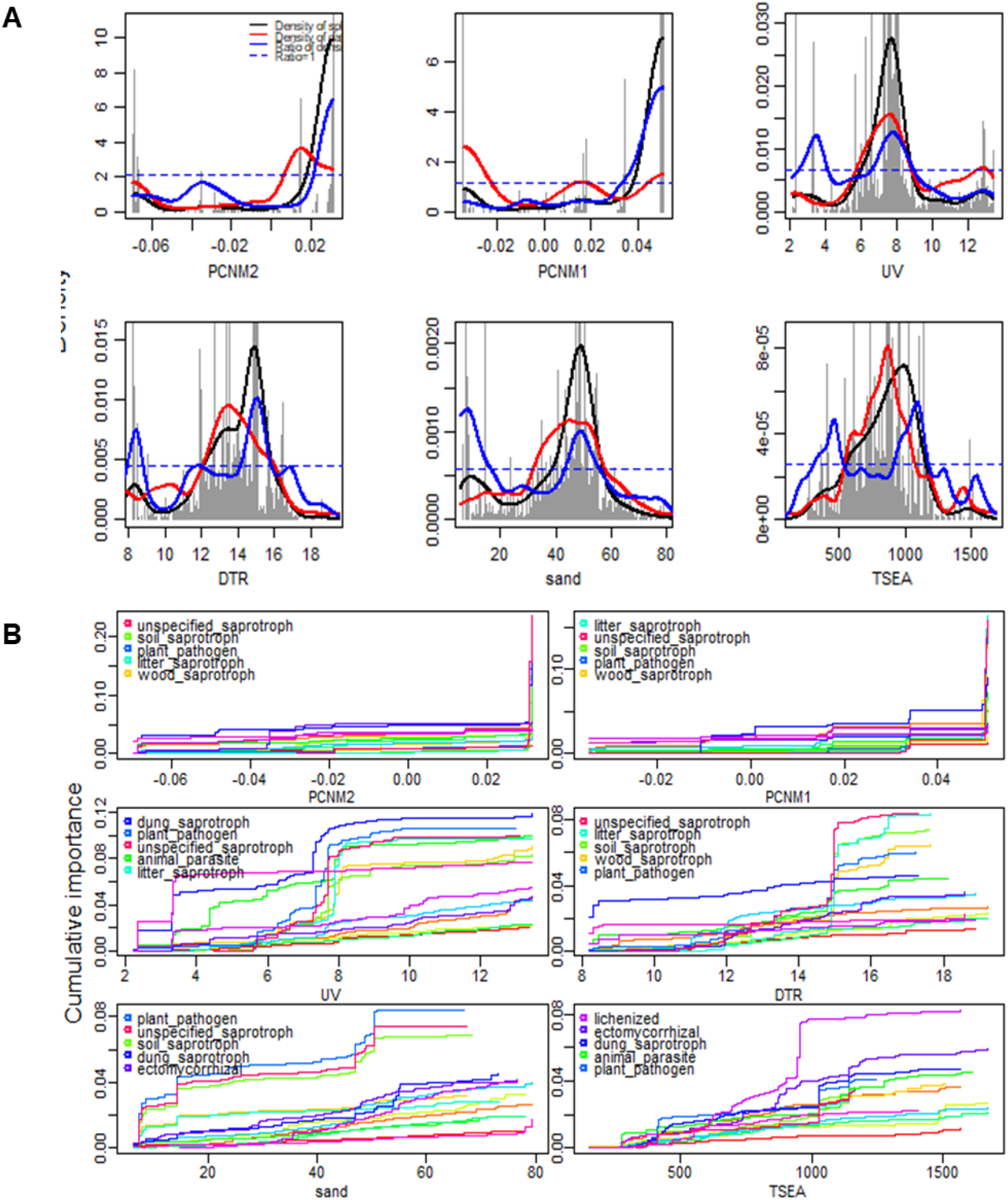
Most relevant predictors of fungal composition in drylands worldwide. **(A)** Frequency histograms of gradient values at which splits occur in the regression trees of the top 15 ecological groups in relationship to the top six environmental variables, showing where along these gradients important compositional changes are taking place. Black lines are the kernel density of the histograms, red lines show the (normalized) distribution of the data along the environmental gradients, and blue lines indicate the ratio between splits and data (ratio between black and red lines). Ratios >1 (above the dotted line) indicate conditions of relatively greater change in genus composition (i.e. community thresholds). Individual plots depict the predictors, arranged (left to right) from the most to the least important. PSEA = precipitation seasonality; TSEA = temperature seasonality; DTR = diurnal temperature range; AI= aridity Index; UV = UV index; MAT = Mean annual temperature. (**B**) Compositional change along the top six environmental gradients for the top five fungal ecological groups. Each line denotes an ecological group and their pattern of compositional change along the gradient. The y-axes have been normalized so that the maximum corresponds to the relative variable importance.

Because the results of the previous analyses do not allow to depict the relationship between environmental predictors and functional guilds of fungi (they only inform about the existence of a high magnitude change affecting the composition of the community), we used Random Forest (RF) models for each guild and SHAP dependence plots (see Online Methods) to visualise these dependencies (Figure 3). All these relationships showed different degree of non-linear behaviours, with marked thresholds in the predictors signalling either abrupt (e.g., changes for DTR, sand, UV or TSEA for unspecified saprotrophs), or non-linear trends (e.g., changes occurring in TSEA) affecting probability of occurrence of fungal ecological guilds. For instance, plant pathogens and unspecified saprotrophs had a higher probability of occurrence with increases in UV (values > 7.5), TSEA (values > 900 SD), and decreases in DTR (values < 14 °C), with pathogens also being positively associated with sand content of approximately 35-50%. Soil saprotrophs were predicted to occur with decreasing UV (values < 7.5), TSEA (values < 500degC), and DTR (values < 14 °C); litter saprotrophs were generally most likely to occur at narrower TSEA (values < 500 SD).

**Figure 3:**
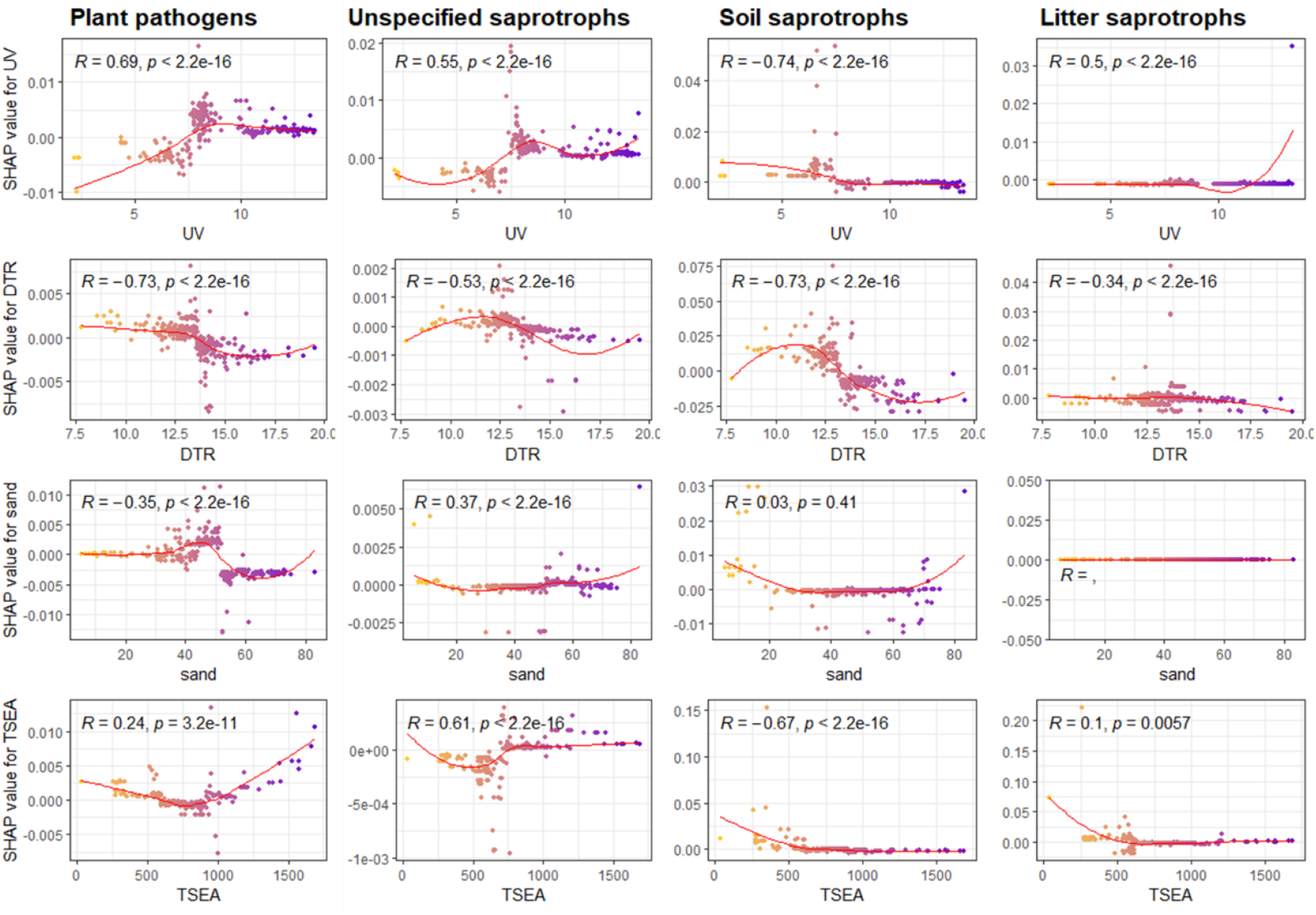
Distribution and environmental predictors of the main fungal ecological groups in drylands. Shapley additive explanations (SHAP) dependence plot of selected climatic and edaphic predictors of plant pathogens and saprotrophs richness in drylands. The effect is expressed as SHAP values, which measure the impact of each predictor on the model output (richness of a particular fungal ecological group). SHAP values are derived for a given predictor value in a process analogous to partial dependence plots; thus each point on the plot corresponds to a prediction in a sample (see methods). *R* = Spearman Rho correlation coefficient; *p* = p value.

### Global patterns of fungal diversity in drylands

The Random Forest model built to assess the relative contribution of environmental predictions of overall fungal richness in drylands revealed a strong contribution of spatial distance (PCNM1) and AI (%IncMSE > 20 for both; Figure 4A). We observed generally similar levels of richness between dry sub-humid, semi-arid and arid biomes, while hyper-arid areas supported a significantly lower (Wilcoxon test, *p* > 0.05) fungal diversity (Figure 4B). Consistently, the fungal maps estimating the expected geographical distribution and richness of dryland fungi (R=0.92, Figure 4C), broadly reflected the extent of well-characterised high classes of aridity, with sharp declines in fungal alpha diversity in hyper-arid regions of the globe.

**Figure 4.**
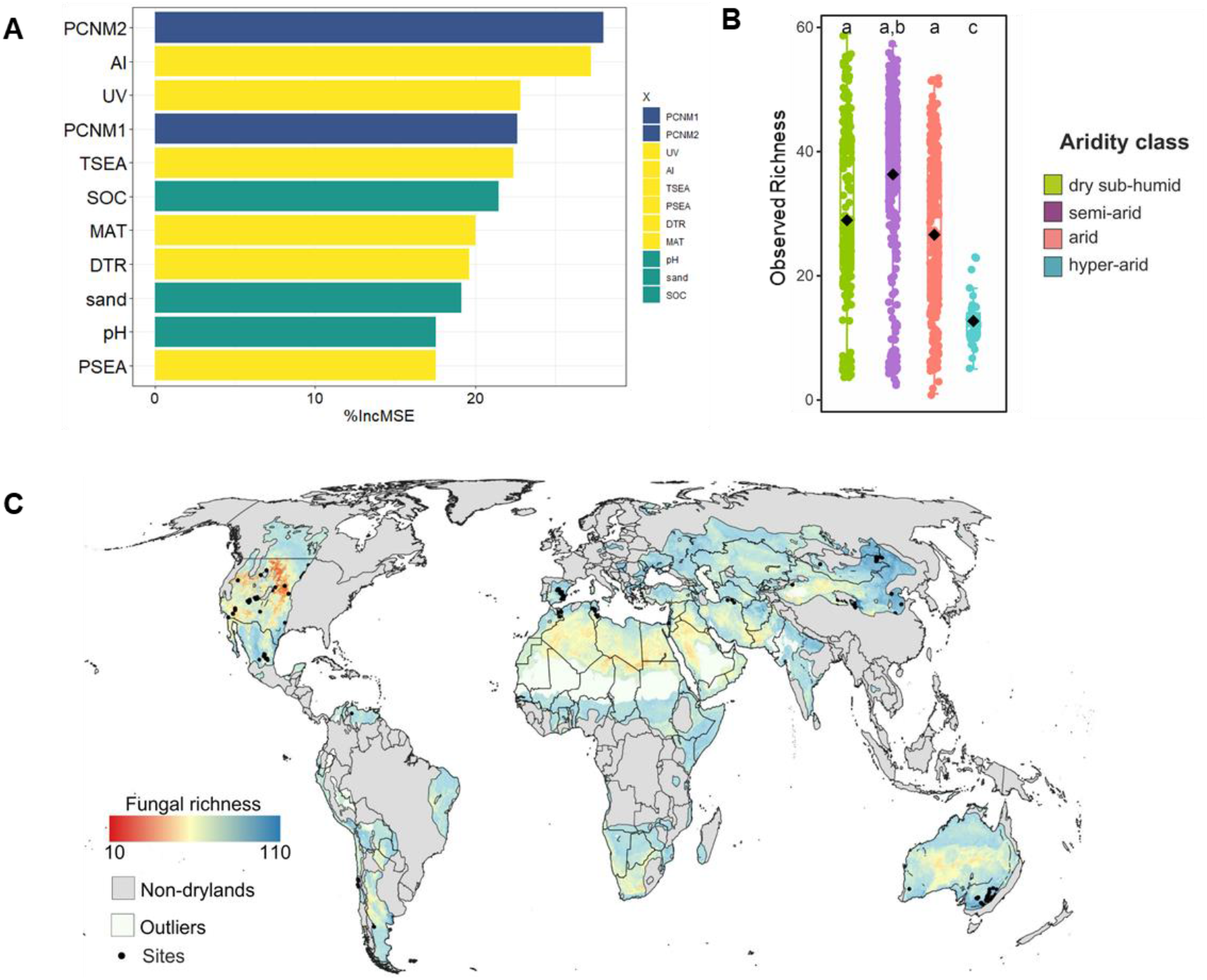
Environmental predictors of dryland fungal community richness. **(A)** Relative importance, expressed as %IncMSE, of each environmental predictor included in the random forest analysis. **(B)** Box-plots illustrating alpha diversity indices (Observed richness) of fungal phylotypes (genus level) for the different aridity classes. Individual data points, median values and interquartile ranges are shown. Different letters indicate significant differences (P < 0.05, Wilcoxon test). (**C**) Predicted global distribution of fungal richness across drylands worldwide. The scale bar represents the observed richness of each ecological group.

## Discussion

Our study demonstrates, for the first time, that environmental gradients related with solar UV radiation (i.e., the UV index), climate seasonality (i.e., DTR and TSEA) and soil structure (i.e., sand content) are critical predictors of fungal community composition in global dryland soils, with the greatest influence detected in association with the richness of putative plant pathogens and a range of fungal saprotrophic groups. Most importantly, we found that the relationships of these environmental predictors with different fungal ecological groups are markedly non-lineal, exhibiting thresholds in the values of environmental variables that may signal particularly vulnerable environmental scenarios. In particular, increases in UV and temperature seasonality above a certain threshold (7.5 and 900 SD, respectively) were associated with an increased probability of plant pathogens and unspecified saprotrophs occurrence, with plant pathogens also being positively associated with sand content of approximately 35-50%. Conversely, these parameters had an overall negative relationship with litter and soil saprotrophs richness, the latter being negatively influenced also by increases in DTR (values > 14 °C).

These trends can be explained by the unique abiotic features that regulate biogeochemical cycling in drylands and the peculiar physiological attributes of different saprotrophic and pathogenic fungal groups ^18^. In most arid lands, temperature-related variables and soil structure are considered critical factors in determining decomposers composition ^19^, and traditional models identify extreme temperatures and low soil moisture typical of dry regions of the world as main controllers of litter quality and microbial activities ^20^. These environmental parameters can thus act as limiting factors for the distribution of fungal decomposer, which tend withstand overall lower temperature ranges compared to other guilds, such as pathogenic fungi ^28^, thus explaining their decrease in occurrence probability with increases in temperature ranges and variability ^15^. Soil structure attributes are also expected to exert various influences on fungal communities, for example by enhancing substrate availability from SOC pools, while also controlling water holding capacity ^21^, which can in turn regulate fungal saprotroph richness and composition.

Similarly, in many arid ecosystems, solar radiation is considered a primary driver of decomposition and carbon cycling ^22,23^, resulting in a significant photo priming effect that controls root exudation, litter quality and nutrient availability, and accelerates abiotic-driven decomposition in these systems ^24^.The tight link between UV radiation and biogeochemical cycling in drylands could thus explain the prominent role of the UV index in predicting the distributions of saprotrophic groups associated with soil and litter, and the overall negative influence on soil saprotroph richness.

Conversely, putative plant pathogens were mostly dominated by ascomycetes from the classes Dothideomycetes and Leotiomycetes, which are known to possess unique physiological traits allowing them to resist environmental stresses typical of drylands, including UV radiation, high temperature fluctuations and dissection ^7,9,25^. Such common traits might allow potential pathogens from these taxonomic groups to adapt to a wide range of environmental stressors, possibly explaining the ability of the members of this ecological group to thrive in extreme environments. Additionally, photoreception and light-dependent traits have been recently suggested as a likely mechanism allowing foliar pathogens from sun-lit habitats to recognize potential partners and stressed hosts ^26^, indicating that increases in UV radiation might have an important but underestimated role in facilitating the establishment of pathotrophic fungi in dryland ecosystems. Our analyses indicate that countries crossing a 7.5 UV radiation index threshold and experiencing high temperature seasonality (values > 900 SD) could be at increased risk of pathogen outbreaks, with potentially detrimental ecological and economic implications.

Overall, the relative importance of the environmental predictors identified in our survey is markedly different from the findings from previous global studies conducted in more mesic environments, where mean annual precipitation has the strongest influence on the richness of most fungal taxonomic and functional guilds^27,28^, reflecting the peculiarity of the environmental attributes that regulate ecosystem functionality in drylands. Interestingly, the aridity index, which is considered a primary driver of change in drylands ^3^, had a secondary role in determining fungal functional changes in our dataset. However, in line with other global surveys, we observed significant decline in fungal alpha diversity with increasing aridity, confirming the critical role of this climatic variable in shaping microbial biodiversity in global drylands. The compositional turnover of the dryland functional mycobiome was also strongly associated with the eigenvector-based spatial descriptors (PCNMs), which were also significantly correlated to the total fungal community richness. At the guild level, the strongest effect was recorded for phototrophic and saprotrophic fungi, the most abundant members of the community in our dataset. The large predictive power of PCNMs could indicate a role for neutral processes, such as dispersion limitation and/or stochastic events, in shaping the community dynamics of the dominant fraction of the fungal assemblies ^29^. Indeed, abundant microbial taxa with higher dispersal rates tend to be affected by drift or priority effects more than their rarer counterparts ^30^, possibly explaining the large influence of spatial variables observed in this study for the most frequent functional groups. However, the overall performance of our models remained relatively low for the less common functional guilds in the dataset, suggesting that other processes such as biotic interactions ^31^, or unmeasured environmental gradients (e.g., vegetation composition ^32,33^) might play a critical role in characterising the distribution of these lower abundance community members in dry ecosystems, and warrant further investigations on a global scale.

Collectively, our results indicate that solar UV radiation, temperature and precipitation variability, and soil structure might be underappreciated drivers of global distribution of critically important fungal groups, such as plant pathogens and saprobes. The relationships between functional composition and environment uncovered in this study are crucial for developing accurate mechanistic models and making predictions about climate-driven shifts in fungal community structure, and thus ecosystem functions. Overall, our findings imply that processes leading to shifts in solar radiation (e.g., stratospheric ozone depletion), soil structure (e.g., land-use change, and land degradation), and seasonal climatic patterns (e.g., increases in atmospheric levels of a greenhouse gases), might have disproportionate consequences for the distribution of fungal groups linked to vegetation and biogeochemical cycling in drylands, and could influence the balance of plant–soil interactions in drylands. These processes might be particularly exacerbated in the Southern Hemisphere, where climate change has profound influence on the ozone layer ^34^, and could be further compounded by predicted increases in extreme heatwave events, which can synergistically alter the UV-mediated effects on terrestrial ecosystems ^35,36^. In particular, we observed a significant threshold in composition turnover at UV index > 7 and diurnal temperature ranges > 900 SD, suggesting that the strongest effects of climate-driven shifts in UV incidence and climate seasonality could occur in regions approaching these values, such as arid regions of the Australia, Centre and South America, North Africa and Central Asia ^37^, with unknown ecosystem-level implications.

Taken together, the comprehensive catalogue of ecology–climate relationships we provide paves the way to a more exhaustive and detailed understanding of the complex role of climate and soil in regulating fungal biogeography, especially in those regions of the world that are most vulnerable to environmental changes, such as global drylands. This work opens a new line of investigation to include quantifying the importance of abiotic and biotic processes that govern fungal communities across contrasting regions of the world, with particular emphasis on identifying the traits, and traits trade-offs, underpinning their functional capabilities in such unique ecosystems ^38^. In particular, we anticipate that as strain-specific trait data become available, better assessment of functional variation expressed within and among communities in relation to UV tolerance and climate variability will be possible. This information is required to provide better predictions of the current and future adaptation of fungi to the effects of climate change, and their ramifications for sustainability of dryland ecosystems.

## Online Methods

### Literature and environmental variables selection

We have undertaken a comprehensive meta-study of data published on the composition of soil fungal communities in drylands across the world. This approach enabled us to re-analyse multiple datasets from different biogeographical regions and biomes and compile a large dataset of fungal taxa distribution worldwide (see Supplementary Material for details on literature selection, bioinformatics data processing, and functional group assignments). In total, 14 studies, encompassing over 912 top-soil (5-10 cm depth) sampling points, were identified and included in the analysis; this allowed us to encompass all continents (including Antarctica; Supporting Information Figure S1), spanning a wide range of environmental conditions. The final sample list included all drylands subtypes (hyper-arid, AI 0.0–0.05, *n* = 42; arid, AI = 0.05–0.20, *n* = 274; semiarid, AI = 0.20–0.50; *n* = 336; dry-sub humid, AI = 0.50–0.65; *n* = 264).

Metadata were collected from the published papers and/or public repositories where they were submitted by the authors, while in a few cases from the authors of individual studies upon request and are included in Supporting Information. Additional metadata were collected from the Worldclim database (https://www.worldclim.org; ~1 km resolution) ^39^, and included spatial, climatic and edaphic parameters. Climatic data included a range of variables related to temperature and precipitation variability that are considered important drivers of fungal distribution at large scales (^28^ – i.e., mean annual temperature (MAT), precipitation seasonality (PSEA), temperature seasonality (TSEA) - as well as the aridity index (AI). The aridity index was obtained from the global maps of ^40^, which provides the averaged AI of the period 1970-2000, and has a spatial resolution of 30 arc-seconds. We also collected data on the AI from the Global Potential Evapotranspiration database ^41^, which is based on interpolations provided by WorldClim. We used aridity index instead of mean annual precipitation in our study because aridity includes both mean annual precipitation and potential evapotranspiration, and is therefore a more accurate metric of the long-term water availability at each site; moreover aridity index is the one used for categorizing drylands and is the one used on global reports about desertification and climate change. UV radiation (UV) was further included given its importance in driving biogeochemical processes in dryland soils ^18,41^. Three important edaphic determinants of fungal biogeography (i.e., % of sand, SOC and pH), obtained from SoilGrids v2 database, were also included, allowing us to evaluate the importance of soil physico-chemical attributes for fungal distribution in drylands. To accommodate spatial variables, principal coordinates of neighbour matrices (PCNMs) were also included as explanatory variables in downstream analyses to examine the importance of spatial filters on community composition ^42^. PCNMs were calculated with the vegan R package, and the first two of the positive PCNMs were retained. We obtained complete environmental metadata for a total of 743 samples, which were used for the quantification of the functional turnover across environmental gradients (see methods below). Downstream analysis, unless otherwise specified, were performed in R environment v. 4 and using the genus-level taxonomy table.

### Quantification of ecological turnover and thresholds across environmental gradients

To explore the environmental drivers of distributions of the most common fungal ecological guilds in global drylands, we modelled their occurrence using an approach similar to ^43^. Briefly, we first identified the most common guilds among those occurring in at least 10% of the samples, which resulted in 16 ecological groups. We then explored the most important environmental predictors of fungal ecological turnover by generating a random forest fitting a total of 500 trees using the extended modelling procedure available in R package “gradientForest” ^44^. The gradient forest (GF) technique derives monotonic, nonlinear functions that characterize compositional shifts along each fitted environmental gradient, without a priori distributional assumptions about the frequency of response variables, as opposed to other methods such as generalized linear models or generalized additive models. The importance of each predictor variable (measured as R-squared) in the model is assessed by quantifying the decrease in performance when each predictor variable is randomly permuted, using a conditional approach which accounts for collinearity between predictor variables ^45^. This allows us to assess each predictor’s importance relative to one another in terms of their influence on patterns of composition. Additionally, GF allows the development of empirical distributions that represent species (ecological groups in this case) turnover along each environmental gradient, by aggregating the values of the tree splits from the Random Forest models for all individual models with positive fits (R-squared > 0). The turnover function is measured in dimensionless R-squared units where groups with highly predictive random forest models (i.e., high R-squared values) have greater influence on the turnover functions than those with low predictive power (i.e., lower R-squared). These turnover functions can provide unique insights into the nature of how functional patterns vary along multiple environmental gradients, at the level of individual ecological groups as well as the mycobiome as a whole when these individual curves are averaged to obtain a global R-squared value. The incremental approach to model fitting in GF makes it also well suited for the analysis of large datasets, whose size can be limiting in other approaches, such as generalized dissimilarity modelling ^46^. A detailed description of these methods can be found in ^44–47^.

Following the GF approach described above, the model performance was assessed by the overall goodness-of-fit (R-squared) of predicted against observed values and by the cross-validated out-of-bag R-squared values per ecological group, while the significance of each environmental variable was assessed by the relative importance weighted by R-squared values ^44^. Subsequently, to visualize the importance and abruptness of specific thresholds and to identify common threshold locations among ecological groups, we plotted their cumulative importance, whereby the shape of the resulting distribution curves describes the magnitude of compositional change along the most important gradients, with the standardized ratio of split density >1 indicating the highest manifestation of a threshold ^48^. The concept of community threshold used here is defined as a zone along an environmental gradient where the change in community composition is enhanced as a result of sharp increases or decreases in the occurrence of several functional groups (depending on the direction of the gradient). Therefore, GF enabled us to identify critical values along environmental gradients that correspond to threshold changes in functional composition.

Finally, to further illustrate the directionality of the response to environmental predictors, we run GradientBoost (GB) models with SHapley Additive exPlanations (SHAP) dependence plots. GB models were run individually for each ecological group and were done solely for the most important environmental variables and ecological groups best explained by the GF models. The SHAP method is derived from game theory and measures how much each feature of a model contributes to the increase or decrease of the probability of a single output with respect to the average of the ones used to train the model (ie, the richness of a particular ecological group in this case). In a nutshell, the SHAP value is derived from a regression tree model for a given feature and prediction. Its value is the effect of the predictor of interest in the model output for a given prediction and, thus it is provided in the same units as the response variable. SHAP values are actually homologous to evaluating the expression β1*x1 in a regular multiple regression (y~β1*x1+β2*x2+…+βn*xn). This means that, essentially, a given predicted value of the model is the summation of all SHAP values obtained from the model given the values of predictors (74). By plotting the values of predictor vs the associated SHAP values we obtain a response curve analogous to the effects of that predictor over the response variable (i.e., a partial dependence plot). SHAP values are widely used in machine learning (75), economics (72), security (73) and ecology (76). SHAP values can be positive or negative, whereby a positive trend indicates that a feature is expected to positively influence the occurrence of a particular ecological group, and vice-versa. Models were built with the “xgboost” package and SHAP values were extracted with the “SHAPforxgboost” package in R.

### Quantification of biodiversity and environmental drivers of fungal community composition

Alpha diversity was estimated using the R package ‘phyloseq’ ^49^, calculating biodiversity indices as the species richness as a count of the observed taxa in each sample. A Random Forest (RF) model was built using the ‘randomForest package with 500 trees in R to assess the relative contribution of climatic, spatial and edaphic predictors on dryland fungal richness. Statistical analysis was performed to identify how overall richness changed across dryland types by one-way analysis of variance (one-way ANOVA) and pairwise multiple comparison procedure (Tukey test); a small probability p-value (<0.05) indicated a significant difference. The extent of the global distribution of soil fungal richness was estimated using a Random Forest regression analysis as described in the supplementary material.

## Data availability

Metadata and samples ID are freely available in Figshare (https://doi.org/10.6084/m9.figshare.19243749.v1)

## Code availability

The codes for the computational analyses are available in Figshare (https://doi.org/10.6084/m9.figshare.19243749.v1).

## Acknowledgements

C.C. acknowledge funding from the Italian National Program for Antarctic Research (PNRA) and is supported by a PNRA postdoctoral fellowship. E.E. is supported by an Australian Research Council DECRA fellowship (DE210101822). M.D-B. is supported by a project from the Spanish Ministry of Science and Innovation (PID2020-115813RA-I00), and a project PAIDI 2020 from the Junta de Andalucía (P20_00879). Microbial distribution and colonization research in B.K.S.’s lab is funded by the Australian Research Council (DP190103714). E.G. is supported by the European Research Council Grant agreement 647038 (BIODESERT).

## Authors’ contributions

E.E. and C.C. developed the original idea of the analyses presented in the manuscript. Literature selection and raw data retrieving were done by C.C. Bioinformatic analyses were done by D.A and C.C.. Statistical analyses, environmental modelling and data interpretations were done by E.E., M.B., E.G., M.D-B., and C.C. The manuscript was written by E.E. with contributions from all the co-authors. All authors have read and agreed to the published version of the manuscript.

